# Neurofeedback linked suppression of subthalamic beta oscillations speeds up movement initialisation in Parkinsonian Patients

**DOI:** 10.1101/687582

**Authors:** Shenghong He, Abteen Mostofi, Emilie Syed, Flavie Torrecillos, Gerd Tinkhauser, Petra Fischer, Alek Pogsyan, Harutomo Hasegawa, Yuanqing Li, Keyoumars Ashkan, Erlick Pereira, Huiling Tan

**Affiliations:** MRC Brain Network Dynamics Unit at the University of Oxford, United Kingdom; Nuffield Department of Clinical Neurosciences, University of Oxford, United Kingdom; Neurosurgery and Consultant Neurosurgeon St George’s University Hospital, London, United Kingdom; Department of Neurology, Bern University Hospital and University of Bern, Bern, Switzerland; Department of Neurosurgery, King’s College Hospital NHS Foundation Trust, King’s Health Partners, London, United Kingdom; School of Automation Science and Engineering, South China University of Technology, Guangzhou, China

**Keywords:** Beta oscillations, neurofeedback training, beta burst, local field potential (LFP), Parkinson’s disease (PD)

## Abstract

Enhanced beta oscillations (13-30 Hz) in the subthalamic nucleus (STN) have been associated with clinical impairment in Parkinson’s disease (PD), such as rigidity and slowing of movement, with the suppression of STN beta activity through medication or deep brain stimulation correlating with improvement in these symptoms. Recent studies have also emphasized the importance of the time dynamics of the STN beta oscillations in the pathology of PD. An increased probability of prolonged beta bursts, defined as periods when beta band power exceeds a certain threshold, was more closely related to motor symptoms than average power; and the occurrence of beta bursts just before a go cue slows cued movements. Here we adopted a sequential neurofeedback-behaviour task paradigm to investigate whether patients with PD can learn to suppress pathological beta oscillations recorded from STN with neurofeedback training and whether the training improves the motor performance. Results from twelve patients showed that, compared with the control condition, the neurofeedback training led to reduced incidence and duration of beta bursts in the STN local field potential (LFP) and also reduced the synchrony between the STN LFP and cortical activities measured through EEG in the beta frequency band. The changes were accompanied by a reduced reaction time in cued movements. These results suggest that volitional suppression of beta bursts facilitated by neurofeedback training could help improve movement initialisation in Parkinson’s disease.

**Significance Statement:** Our study suggests that a neurofeedback paradigm which focuses on the time dynamics of the target neural signal can facilitate volitional suppression of pathological beta oscillations in the STN in Parkinson’s disease. Neurofeedback training was accompanied by reduced reaction time in cued movements, but associated with increased tremor in tremulous patients. The results strengthen the link between subthalamic beta oscillations and motor impairment, and also suggest that different symptom-specific neural signals could be targeted to improve neuromodulation strategies, either through brain stimulation or neurofeedback training, for patients with tremor and bradykinesia-rigidity.

## Introduction

Enhanced synchronization of neural activity in the beta band (13-30 Hz) has been consistently observed in the subthalamic nucleus (STN) in patients with Parkinson’s disease (PD), and correlates with motor symptoms of PD such as rigidity and bradykinesia (Kühn et al. 2006, 2009, Little et al. 2012). Treatment of PD, either through dopaminergic medication or deep brain stimulation (DBS), suppresses this pathological synchronization, and the degree of suppression correlates with the improvement in motor symptoms (Jenkinson and Brown, 2011). STN beta activity in PD is not continuously elevated but fluctuates over time. Synchrony in this frequency band takes the form of short-lived bursts of different durations and amplitudes (Tinkhauser et al., 2017a, b). The percentage of longer beta bursts with large amplitude in a given interval positively correlates with clinical impairment in the disease (Tinkhauser et al. 2017a). Closed-loop DBS responsive to the fast temporal dynamics of beta activity limits the evolution of the beta synchrony over time. By selectively truncating long beta bursts, this approach has been reported to achieve a superior clinical effect compared with conventional continuous DBS even though both approaches reduce average β-band power (Little et al. 2013, 2016). This highlights the importance of modulating the temporal dynamics of the beta activity in the treatment of Parkinson’s disease.

Neurofeedback training, which aims to train subjects to self-regulate their neural activity, has been proposed to be a promising technique with which to tune pathological brain oscillations corresponding to different diseases (Ros 2014). For example, Subramanian et al. developed a real-time functional magnetic resonance imaging (rtfMRI)-based neurofeedback paradigm and showed that patients with PD were able to alter local brain activities in supplementary motor area (SMA) and improve motor function (Subramanian et al. 2011). Patients with PD can also learn to modulate beta oscillations in the sensorimotor cortex measured using the electroencephalogram (EEG) (Erickson-Davis et al., 2012; Azarpaikan et al., 2014) or an electrocorticography (ECoG) electrode grid placed on the surface of the cortex (Khanna et al. 2017). However, none of these studies showed any motor improvement associated with the neurofeedback training. This may be due to the fact that cortical β-power may not be a good biomarker for Parkinsonian symptoms since cortical beta power is similar in PD patients with and without DBS or levodopa therapy (de Hemptinne et al. 2013, Rowland et al. 2015) or even in PD and non-PD patients (de Hemptinne et al. 2015). PD patients were also shown to be able to voluntarily control the β-band power in STN measured using electrodes implanted for deep brain stimulation in the instructed direction after 10 minutes of continuous neurofeedback training (Fukuma et al. 2018). However, behavioural improvement was not demonstrated in these patients.

In this study, we adopted a sequential neurofeedback-behaviour task design of the kind that has helped to shed light on the relationship between neural activity and behaviour (McFarland et al., 2015; Khanna and Carmena, 2017). In the current paradigm, the neurofeedback targets the incidence of high amplitude beta bursts, and the neurofeedback training was followed by a cued finger pinch movement. Results from twelve participants with PD showed that training using this paradigm allowed the patients to learn to not only modulate the average beta power, but also modulate the temporal dynamics of beta oscillations in STN LFPs in the desired direction with reduced incidence of beta bursts per unit time and reduced average beta burst duration within STN. The training also led to reduced synchrony between STN and ipsilateral motor cortex. These changes were specific to the targeted β-band only, and were associated with reduced reaction time in subsequently cued movements. Our results suggest that neurofeedback training may be a promising technique to improve motor initialisation in PD, and strengthen the link between beta, particularly beta bursts, and motor impairment.

## Materials and Methods

### Subjects

Twelve PD patients (21 hemispheres, 4 females) who underwent bilateral implantation of DBS electrodes targeting the motor area of the STN participated in this study. All patients responded to dopaminergic medication evaluated using Unified Parkinson’s Disease Rating Scale (UPDRS), and had normal or corrected-to-normal vision. All experiments were conducted with the patients off any dopaminergic medication. The study was approved by the local ethics committees and all patients provided their informed written consents according to Declaration of Helsinki before the experiments.

### Experimental protocol

Our neurofeedback training protocol comprised of multiple short trials (shown in Fig. 1). Each trial consists of a 2-s period where the patients are instructed to get ready, a neurofeedback phase lasting 4 to 8 seconds, followed by a movement go cue. During the neurofeedback phase of each individual trial, a basketball appears in the top-left corner of the screen and moves toward the right at a fixed horizontal speed. Meanwhile, the vertical movement of the basketball is sensitive to the STN beta power calculated in real-time. For each update, if the calculated beta power (shown as the height of the black bars in Fig. 1A) is larger than a predefined threshold *T* (shown as the red line in Fig. 1A), the basketball will drop downwards by a fixed distance; otherwise, the ball will keep its height on the screen. Thus, the vertical position of the basketball indicates the incidence of high amplitude beta bursts, but the participants were blind to the other variations in the beta power lower than the threshold value. This design reduces the noise in the visual feedback that is not related to the pathology of Parkinson’s disease, with the aim of helping the participants to gain a sense agency within a short time. In the neurofeedback training condition, participants were instructed to try to keep the ball floating on the top until the end of the basketball presentation. The patients were explicitly told that imagining their contralateral hand movements may help to improve the performance but were encouraged to try different strategies as well. In order to control for the effects of the moving visual stimuli and attention, participants also performed the task in a ‘no-training’ condition, in which the participants were instructed to pay attention to the ball movement and get ready for the go cue without having to control the position of the ball.

**Figure 1.**
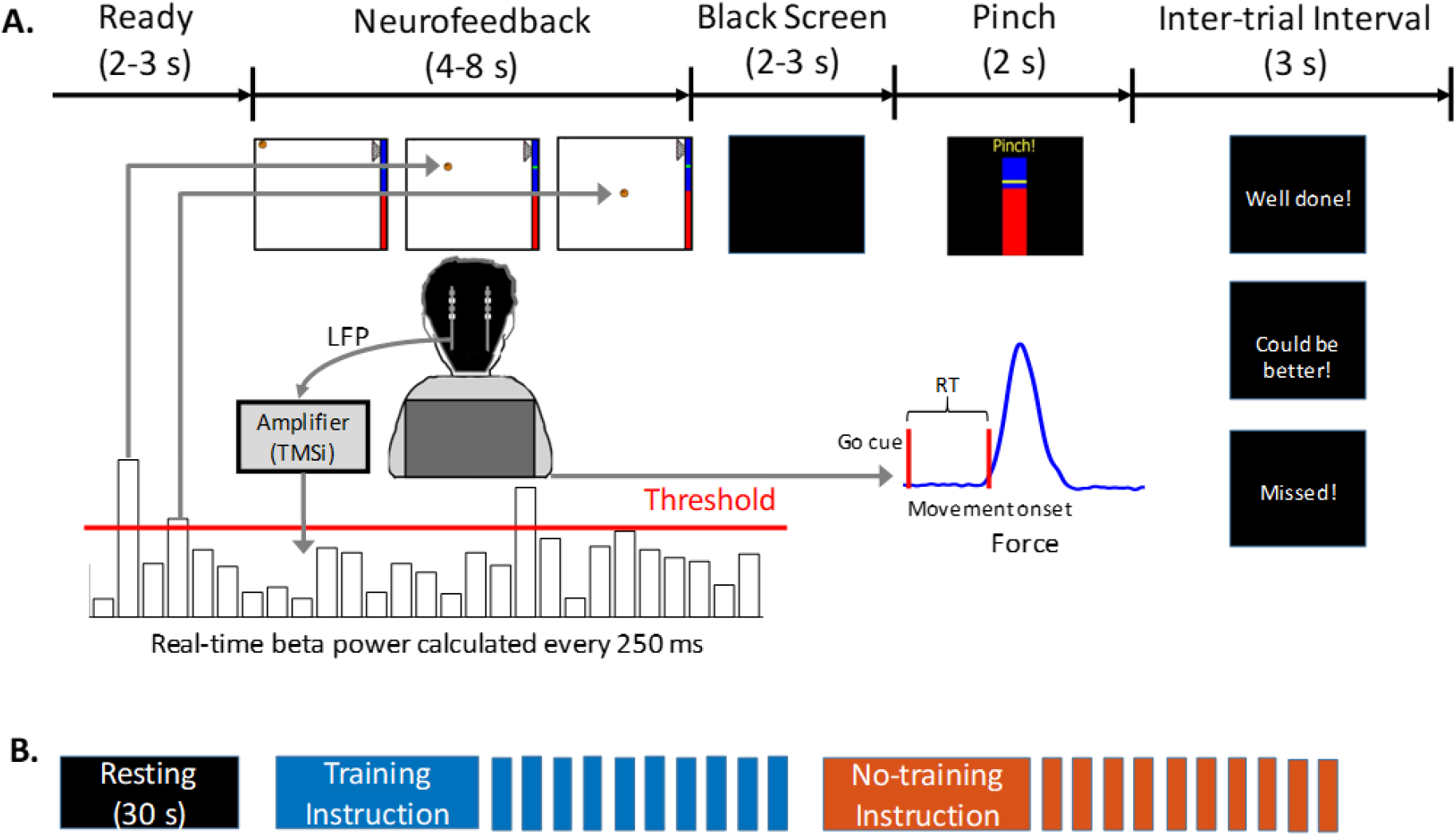
Experimental protocol. (A) Time line of one individual trial. (B) Time line of one training session, the order of the training and no-training block was randomised for each session.

The participants were instructed to perform a thumb finger pinch movement as fast as possible in response to the Go cue to generate a force overshooting a predefined comfortable force level, i.e., 50% of the maximum force (shown as the yellow line in Fig. 1A). All participants were reminded to avoid any voluntary movements until the Go cue was presented, and then to pinch the force meter as fast as possible in response to the Go cue. Feedback about the pinch movement was also provided to the participants at the end of the trial in order to engage the participants and to encourage them to be as fast as possible throughout the experiment. A message ‘Well done!’ or ‘Could be better’ was displayed depending on whether the reaction time of the previous movement was shorter or longer than 800 ms. If movement onset was not detected within 2 s after the Go cue, the message ‘Missed!’ was displayed.

Each experimental session consists of 30 seconds of rest, a ‘training’ block of 10 continuous trials of neurofeedback, and a ‘No-training’ block of 10 continuous trials in the ‘no-training’ condition (shown in Fig. 1B). The order of training and no-training blocks was randomized in each session. Each training/no-training block started with a 10-s duration instruction specifying the requirement of the current block and followed by 10 trials of training/no-training. During the 30-s rest period, power of the selected beta frequency was calculated every 250 ms in order to determine the threshold *T* for triggering the vertical movement of the basketball in the current session (details will be described later). In this study, the beta power threshold and the distance of each drop of the basketball were set so that, if the patient was in resting state, the basketball would drop down to the bottom of the screen within 4-8 s due to the spontaneous variation in the power of beta oscillations found in the rest period. Breaks were provided between sessions and the recording for each STN lasted for around 30 minutes. All participants completed 4 sessions of the task with the dominant hand for the motor task and contralateral STN for neurofeedback. Nine participants performed the task with the non-dominant hand and contralateral STN as well, resulting in 21 data sets in total for Day 1. Four out of the twelve patients repeated the same task over two consecutive days with both sides, which allowed us to investigate cross-day learning effects.

### Data recording

All recordings in this study were undertaken 3-6 days after the first surgery for bilateral DBS electrode (quadripolar macroelectrode, model 3389, Medtronic) implantation and prior to the second surgery for connecting the electrodes to the subcutaneous pulse generator. Eight monopolar channels of bilateral STN LFPs and eight monopolar channels of EEG signals covering “Fz”, “FCz”, “Cz”, “Oz”, “C3”, “C4”, “CP3”, and “CP4” according to the standard 10-20 system, were recorded using a TMSi Porti amplifier (TMS International, Netherlands) at a sampling rate of 2048 Hz. The ground electrode was placed on the left forearm. A common average reference was applied to all monopolar signals. Electromyography (EMG) was simultaneously recorded using the same amplifier from Flexor Carpi Radialis of both arms and the masseter muscle. One tri-axial accelerometer was taped to the back of each hand in order to monitor movements and any tremor. Generated force in the cued pinch movements was recorded using a pinch meter (Biometrics Ltd). In addition, the real-time positions (X, Y) of the basketball in each trial, which allowed evaluation of the performance of neurofeedback training during the online experiment, and the trigger signals of the paradigm were recorded through an open-source toolkit named Lab Streaming Layer (LSL). The synchronization between different data streams was achieved through LSL and another open-source toolkit named Openvibe. The paradigm used in this study was developed in C++ (Visual Studio 2017, Microsoft) and the online/offline data processing was achieved in Matlab (R2018a, MathWorks, US).

### Selecting the STN LFP channel and the targeted frequency band

Prior to the online experiment, each patient participated in a calibration procedure in order to determine the targeted bipolar LFP channel and the individual specific beta frequency band. Bilateral STN LFPs and EEG data were recorded during 60 seconds of rest and 15 trials of cued finger pinch movements with each hand (similar to those used in Fischer et al. 2017). Then we constructed a bipolar LFP channel using each pair of two spatially adjacent depth electrode contacts (i.e., 0-1, 1-2, and 2-3) which resulted in three bipolar LFP channels for each recorded STN. The continuous wavelet transform was applied to calculate the time-evolving power spectra of each trial by using an open source toolkit named fieldtrip. The average time-evolving power spectra of changes in the selected STN LFP channel induced by movements were very similar to those previously observed in STN (Tan et al. 2016). This identified power increases in the theta band (4–8 Hz), power reductions in the broad beta range (13–34 Hz), and power increases in the gamma band (56-65Hz) during movements. Then we calculated the movement-related power reduction for each bipolar LFP channel contralateral to the performing hand in the beta frequency band (13-30 Hz). Finally, the bipolar LFP channel with maximal movement-related power reduction in the 13-30 frequency band was selected as the target LFP channel. A 5 Hz frequency band around the frequency showing maximal movement-related modulation ([*f-2, f+2*]) was determined as the individual specific beta frequency band. The frequency showing maximal movement-related modulation ranged from 17.4 Hz to 21.4 Hz across all tested hemispheres. The average power spectral density of the selected bipolar channel during rest was also quantified using the Fast Fourier Transformation (FFT). As shown in Fig. 2, an obvious peak was observed in the selected beta band in the selected STN LFP channel during rest. The frequency showing maximal movement-related modulation ([f-2, f+2]) was used as the target frequency in our neurofeedback paradigm.

**Figure 2.**
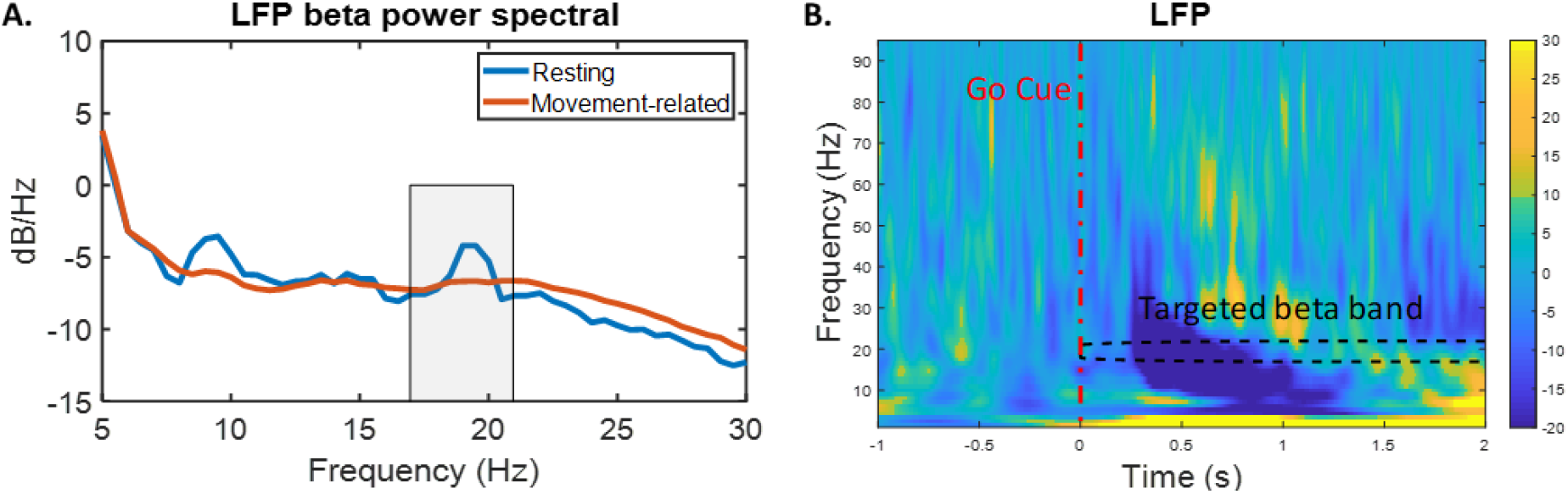
Power spectra of the targeted STN LFP signals averaged across x hemispheres. (A) Resting (blue) and movement-related, non-feedback training (orange) power spectral density (PSD) in STN LFP, rectangle indicates the average of the targeted beta frequency bands. (B) Group average time-frequency power spectra during pinch, red dash line and black dash rectangle indicate the go cue of the pinch and the targeted beta band, respectively. Bar indicates percentage changes relative to baseline.

### Estimating beta power in real-time to determine the position of visual feedback online

During the online experiment, the beta power of the selected frequency band was calculated in real-time every 250 ms using a segment of 500 ms data (with 50% overlapping) recorded from the selected bipolar LFP channel. For each segment of 500-ms data, we first applied a mean subtraction followed by a 5-85 Hz band pass filter on the raw data. Next, FFT was applied to calculate the power spectrum of the filtered data and the average power of the selected frequency band was quantified as the beta band power of the current update. At the beginning of each session, data were recorded with the participant resting for 30 seconds, during which time the beta band power was also updated every 250 ms 119 times. From these values, we selected the 75^th^ percentile as the threshold *T* for that recording session, which means that when the patient was at rest, there was 25% of the time when the beta power would exceed the threshold (Tinkhauser et al., 2017a, b). The threshold was re-calculated at the beginning of each session in order to correct for any drifting in the average beta power with time spent in the task.

In this paradigm, the visual feedback was the vertical position of the basketball at it moved on the screen. The position of the basketball was updated every 250 ms, which corresponded to 16-32 updates during the 4-8 s of neurofeedback in each trial. For each update, the displacement of the basketball on the horizontal axis was constant, so the basketball moved from left to right at constant speed. The displacement of the basketball on the vertical axis was related to the beta band power calculated in real-time. When the updated beta power was larger than the threshold *T*, the basketball displayed on the screen dropped downwards by one step. The distance of each step was calibrated so that the basketball would drop to the bottom of the screen if calculated beta was over the threshold for 25% of the update time points during the feedback phase (4 – 8 s). Thus, the final vertical position of the basketball in each trial was directly associated with the number of incidences when beta power exceeded the threshold within that time window.

## Data analysis

### Visual feedback movement

The trajectory of the visual cursor of the neurofeedback – the basketball movement and the final vertical position of the basketball in each individual trial – were recorded. The difference between the final vertical positions of the basketball between the two conditions, training and no-training, indicated the effect of the neurofeedback training. In addition, from the difference in the ball’s final vertical positions between these two conditions across sessions, we determined the learning effect of neurofeedback training across sessions carried out within one day or over separate days.

### Motor performance

We quantified the reaction time (RT) and the rate of force development in response to the Go cue for each trial based on the recorded pinch force, in order to investigate the effect of neurofeedback training on the performance of the motor task. Force measurements from individual trials were visually inspected and those trials with obvious artefacts and those in which the patients failed to pinch within 2 second were excluded. Thus, for each of the 21 STN hemispheres there were 37.0 ± 2.9 and 36.3 ± 4.2 trials in the training and no-training conditions, respectively, resulting in 1541 trials in total across all tested hemispheres. The measured force was first band-pass filtered between 0.5-20 Hz using a 4^th^ order zero-phase digital filter and then segmented into 4 s epochs extending between 1-s prior to and 3-s after the go cue. We then calculated a threshold by taking the mean plus 3 times the standard deviation (SD) of a segment of 500-ms force data before the cue of the pinch task. The time delay between the go cue and the time point when the force crossed the determined threshold was taken as the RT of that trial. Peak force rate was also calculated by taking the maximum of the first-order differentiation of the filtered force.

Meanwhile, hand tremor was monitored by a tri-axial accelerometer attached to the back of each hand. The power in the tremor frequency band (3-7 Hz, Schneider et al., 2007) was quantified separately and then averaged across all axes.

### STN-LFP beta power

The LFPs from the selected STN bipolar channel and EEGs were further analysed off-line with Matlab (v2018a, MathWorks, US). These signals were first band-pass filtered between 0.5-100 Hz and notch filtered at 50 Hz using a 4 order zero-phase digital filter. After that, we segmented these data into 6 s epochs extending from 2 s prior to and 4 s after the onset of the basketball movement. We stopped epochs at 4 s after onset of basketball movement so that we had a similar duration for analysis across all patients and trials. Time-frequency decomposition of individual trial data was obtained by continuous complex Morlet wavelet transformation with a linear frequency scale ranging from 1 Hz to 95 Hz and linearly spaced number (4-8) of cycles across all calculated frequencies. Then, the average power of the selected beta band within 4 s after the onset of the basketball movement was calculated and decibel (dB) transformed for each individual trial. Next, for each hemisphere, the time-frequency power-spectra were separately averaged across trials in the training and no-training conditions and baseline normalized against a 2 s baseline time window before the onset of the basketball movement by calculating the percentage change in power. The beta power time course in the training or no-training condition was estimated by calculating the mean power of the selected beta frequency band in the corresponding condition.

### STN-LFP beta bursts

In order to investigate the impact of neurofeedback training on the characteristics of beta bursts, we quantified the total duration, average duration and number of bursts occurring in the targeted STN LFP channel for each training/no-training trial. The details of quantifying these burst characteristics have previously been described (Tinkhauser et al. 2017a). For offline analysis, the threshold for the selected bipolar STN LFP channel was determined as the 75^th^ percentile of the beta amplitude calculated based on the 30-s of resting data recorded at the beginning of the first experimental session. The beta amplitude calculated for each time point and each individual trial was compared against the threshold, and those periods of time exceeding the beta amplitude threshold for longer than 100 ms were recognized as beta bursts. Then the total duration (d_1_), average duration (d_2_), and number (n) of bursts during the first four seconds of the neurofeedback phase were further computed for each trial. This enabled comparison of burst characteristics between the training and no-training conditions. In addition, in order to investigate whether there would be a similar impact of neurofeedback training on the bursts in other non-targeted frequency bands, we repeated the burst detection procedure in two other frequency bands by shifting the centre frequency band by 8 Hz down and up, to give “beta-8 Hz” and “beta+8 Hz” frequency bands.

### STN-cortical coherence

The coherence between basal ganglia and cortical sites has been suggested to be increased and reflect pathological coupling in PD patients (Kato et al. 2015). Treatment of PD with dopaminergic medication reduces this coupling and improves symptoms (George et al. 2013). In addition, Tinkhauser et al. (2018) found that beta bursts involve long-range coupling between structures in the basal ganglia-cortical network that is greater during long bursts compared to short beta bursts. Therefore, we also investigated the impact of neurofeedback training on the coherence between STN and the motor cortex. To do so, the STN LFPs from the selected bipolar channel and the monopolar EEG over the ipsilateral motor cortex (“C3” or “C4”) were first z-score-normalized and then transformed into the time-frequency domain through complex Morlet wavelet transformation between 1 and 95 Hz with frequency resolution of 1 Hz. The phase synchrony index (*PSI*) and spectral coherence between the targeted bipolar LFP and ipsilateral EEG channel were calculated for each frequency during the neurofeedback phase for each trial and each experimental condition (training or no-training). The phase synchrony index (*PSI*) was calculated by taking the average of the phase angle differences between the selected bipolar LFP and monopolar EEG channels over time (Lachaux et al. 2000), as shown in formula 1. The spectral coherence was calculated by taking the average of magnitude-modulated phase angle differences between these two channels, normalized against by the power (Lachaux et al. 1999), as shown in formula 2:

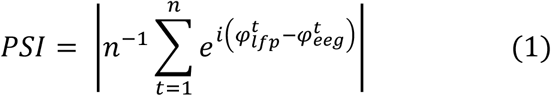

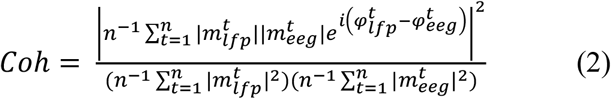

where *n* indicates the total time points in each trial (4 s), 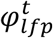 and 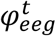 indicate the phase values of the selected LFP and EEG signals at time point 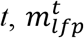 and 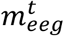 indicate the amplitude values of the selected LFP and EEG signals at time point *t*, respectively.

### Statistical analysis

Two-way ANOVA (using IBM SPSS Statistics 22) with factors of experimental condition (training and no-training) and experimental session (4 in total) was used to evaluate the effect of the neurofeedback training, session number and any potential interaction on neurofeedback task performance in the form of the motor task reaction time or, separately, force. Assumption of sphericity was checked with Mauchly’s test, and if violated F and p values were reported with Greenhouse-Geisser correction. Kruskal-Wallis Test (a non-parametric one-way ANOVA) or paired t-test with Bootstrap (1000 permutations) was used for further post-hoc analysis with Bonferroni correction for multiple comparisons if the data was normally distributed, otherwise, Wilcoxon signed rank test was applied. Mean ± Standard Error, F or t values and associated p values are reported.

Linear mixed effects regression modelling (LME implemented using the Matlab function *fitlme*) was used to assess the within subject relationship between different STN activities and motor performance, and how this was changed by neurofeedback training. In each multilevel linear model, data from all valid individual trials in both experimental conditions (with or without neurofeedback training) from all tested hemispheres were considered. The slope(s) between the predictor(s) and the dependent variable were set to be fixed across all hemispheres; a random intercept was set to vary by hemisphere. LME incorporating data from all individual trials was also used to investigate the effect of cross-day training, in order to increase the statistical power given that fewer patients were recorded over two consecutive days. Significant slopes with estimated p values were reported.

## Results

### Neurofeedback control was achieved within one day of training

Each participant performed the task with each STN and contralateral hand separately for 4 sessions (each containing 10 continuous trials of neurofeedback training and 10 continuous trials of no-training) within one day. LME confirmed strong linear correlations between the final basketball position on the vertical axis and the average beta power (k = −0.047 ± 0.004, p < 0.0001), and with the total time when beta bursts were present during the period visual feedback was presented (k = −0.219 ± 0.020, p< 0.0001). This provides evidence that the final basketball position in the paradigm was influenced by the visual feedback related to beta bursting. A higher final position of the basketball was associated with a reduced average beta power and reduced total duration of beta bursts. The bigger the difference in the final basketball position between the training vs no-training condition the better neurofeedback control of the targeted beta oscillation.

The average final basketball position in the vertical axis was calculated for each experimental session and condition. Two-way ANOVA with factors of experimental condition (‘training’ or ‘no-training’) and experimental session (4 in total) identified significant interaction between experimental session and experimental condition (F_3,60_ = 3.767, p = 0.015) across all tested STNs. This indicated a difference in the effects of neurofeedback training across different experimental sessions. As shown in Fig. 3B, the difference in the final basketball vertical position between the ‘training’ and ‘no-training’ progressively increased from Session 1 to Session 3, suggesting a learning effect within the first 3 sessions, but reduced again in Session 4. Post-hoc analysis using paired t-tests showed that the final basketball vertical position was significantly higher in the ‘training’ condition compared to the ‘no-training’ condition, but only in Session 3 after correction for multiple comparison (t_20_ = 4.096, p = 0.001). It should be noted that the threshold against which real-time calculated beta power was compared was re-calculated for each session in order to control for any potential drift in the beta power. One-way ANOVA applied on the threshold used for different sessions showed that there was no consistent difference in the thresholds used in different sessions (F_3,57_ = 1.494, p = 0.226), suggesting that the difference in the neurofeedback task performance across sessions was not due to changes in the threshold setting. In order to investigate whether the ‘learning effect’ might relate to physical movement during training, we rectified and averaged the amplitudes of all EMG channels across all hemispheres and all trials. As shown in Fig. 3C, EMG activities in training and no-training conditions were very similar, which demonstrated that the learning effect was not consequent on physical movement.

**Figure 3.**
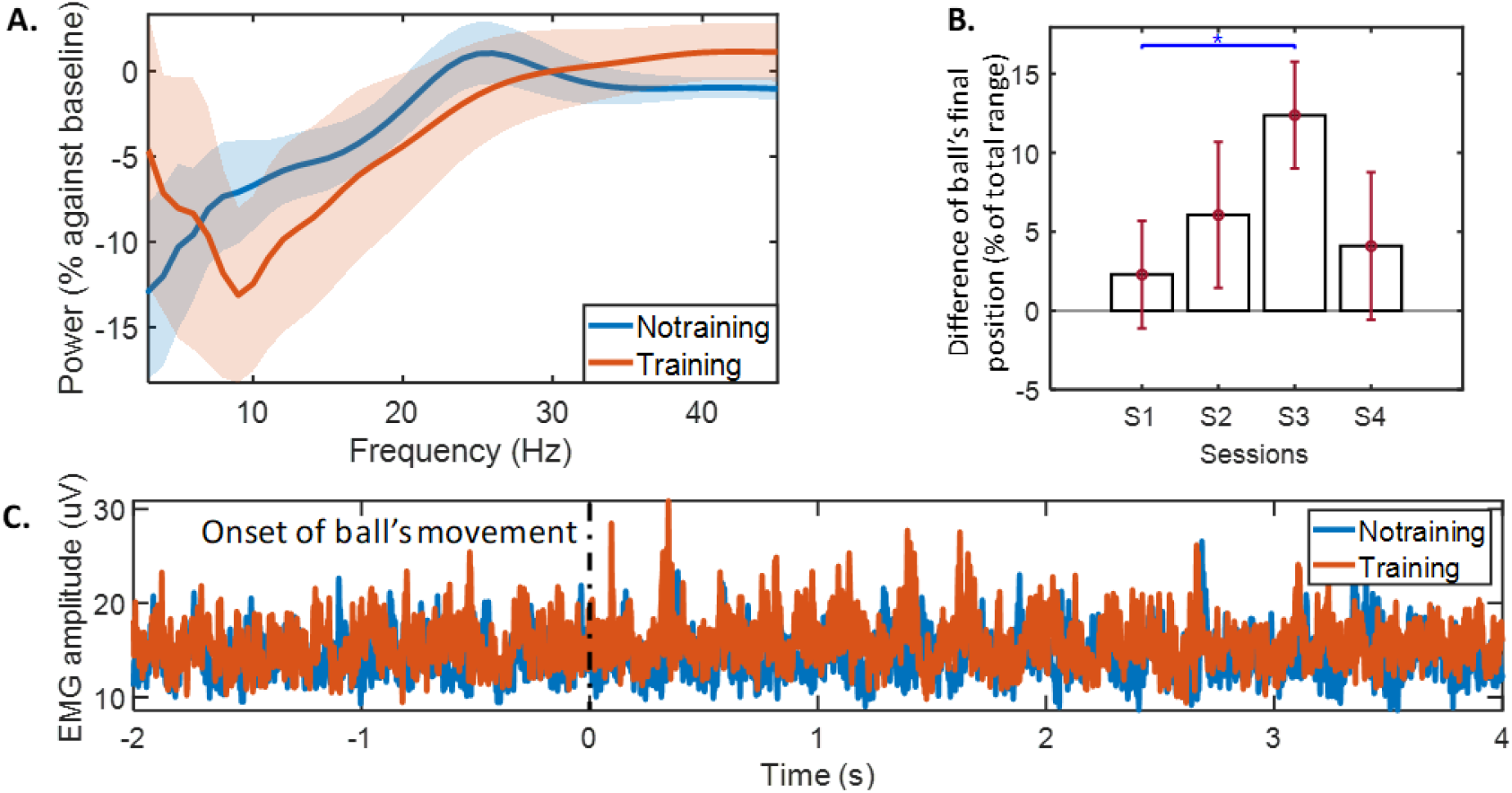
(A) Group average decibel (dB) normalized power spectra across different frequencies during neurofeedback training (orange) or no-training (blue). (B) Group average differences in the ball’s final position between training and no-training conditions across different sessions. (C) Group averaged rectified EMG amplitude in training and no-training conditions, black dash indicates the onset of basketball movement.

### Neurofeedback training led to reduced beta oscillations in STN LFPs

The average time-evolving power spectra of changes in the selected STN LFP channel induced by neurofeedback was derived by aligning the power spectra to the onset of the basketball presentation, normalizing the power of each frequency to the average power of that frequency over the 2 s ‘ready’ time window (Fig. 1) and averaging across all individual trials and across all tested hemispheres. Compared to the ‘ready’ period of time, activity in STN was reduced over a broad frequency band (7–30 Hz) during neurofeedback training (shown in Fig. 3A), similar to the actual movement related modulation shown in Fig. 2B.

Two-way ANOVA was used to investigate the effect of neurofeedback training on the average power of the targeted beta frequency band and the beta burst characteristics during the neurofeedback phase. This confirmed significant effect of neurofeedback in reducing the average beta power (F_1,20_ = 13.01, p = 0.0018) (shown in Fig. 4). The neurofeedback training also reduced beta bursts in the STN LFPs, with reduced percentage of time when the beta amplitude was over the predefined threshold (F_1,20_ = 13.13, p = 0.0017, 16.5 ± 1.6 % compared to 20.5 ± 2.0%) and reduced average burst duration (F_1,20_ = 6.72, p=0.0174, 320.2 ± 26.8 ms compared to 368.9 ± 26.7 ms) (shown in Fig. 5). Even though the average across all tested patients (Fig. 3A) showed a trend of reduction in the activity at the alpha frequency band, this change was not significant across patients (“Beta-8Hz” (centred between 9-12 Hz), average power: F_1,20_ = 0.68, p = 0.4191; total burst duration: F_1,20_ = 0.61, p = 0.4433; average burst duration: F_1,20_ = 0.1, p = 0.7506). There was also no change in the higher frequency band (“Beta+8”, average power: F_1,20_ = 0.35, p = 0.559; total burst duration: F_1,20_ = 0.01, p = 0.9319; average burst duration: F_1,20_ = 0.07, p = 0.8005). These results suggested that during neurofeedback training, the activity in the targeted beta band in the STN LFP was reduced; in addition, beta bursts happened less frequently and were of shorter duration compared with the no-training condition.

**Figure 4.**
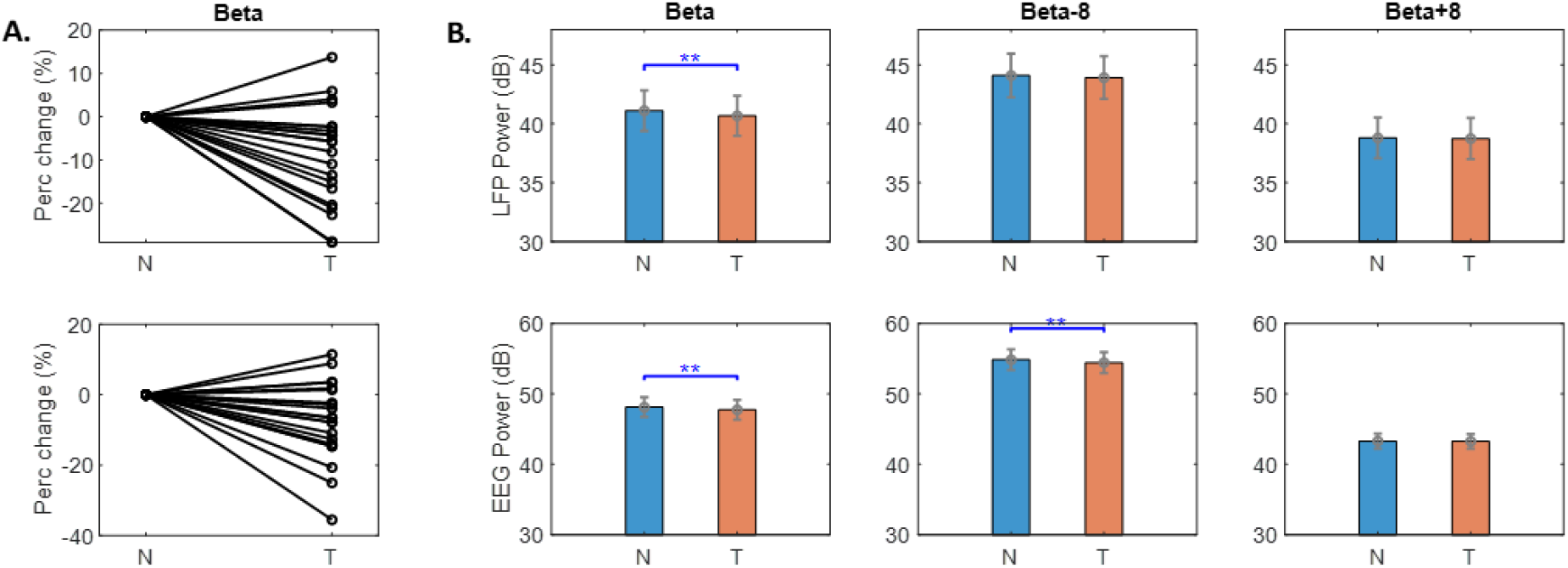
Beta frequency power in LFP and EEG was reduced with neurofeedback training. (A) The percentage change of beta power in LFP (upper) and EEG (lower) for each individual hemisphere. (B) Group average power in beta (left), beta-8 (middle), and beta+8 (right) frequency bands in training (T) and no-training (N) conditions in LFP (upper) and EEG (lower). Values are quantified based on the first 4 s data when the ball was shown and presented as mean ± SEM; **p<0.01; Beta indicates person specific beta band.

**Figure 5.**
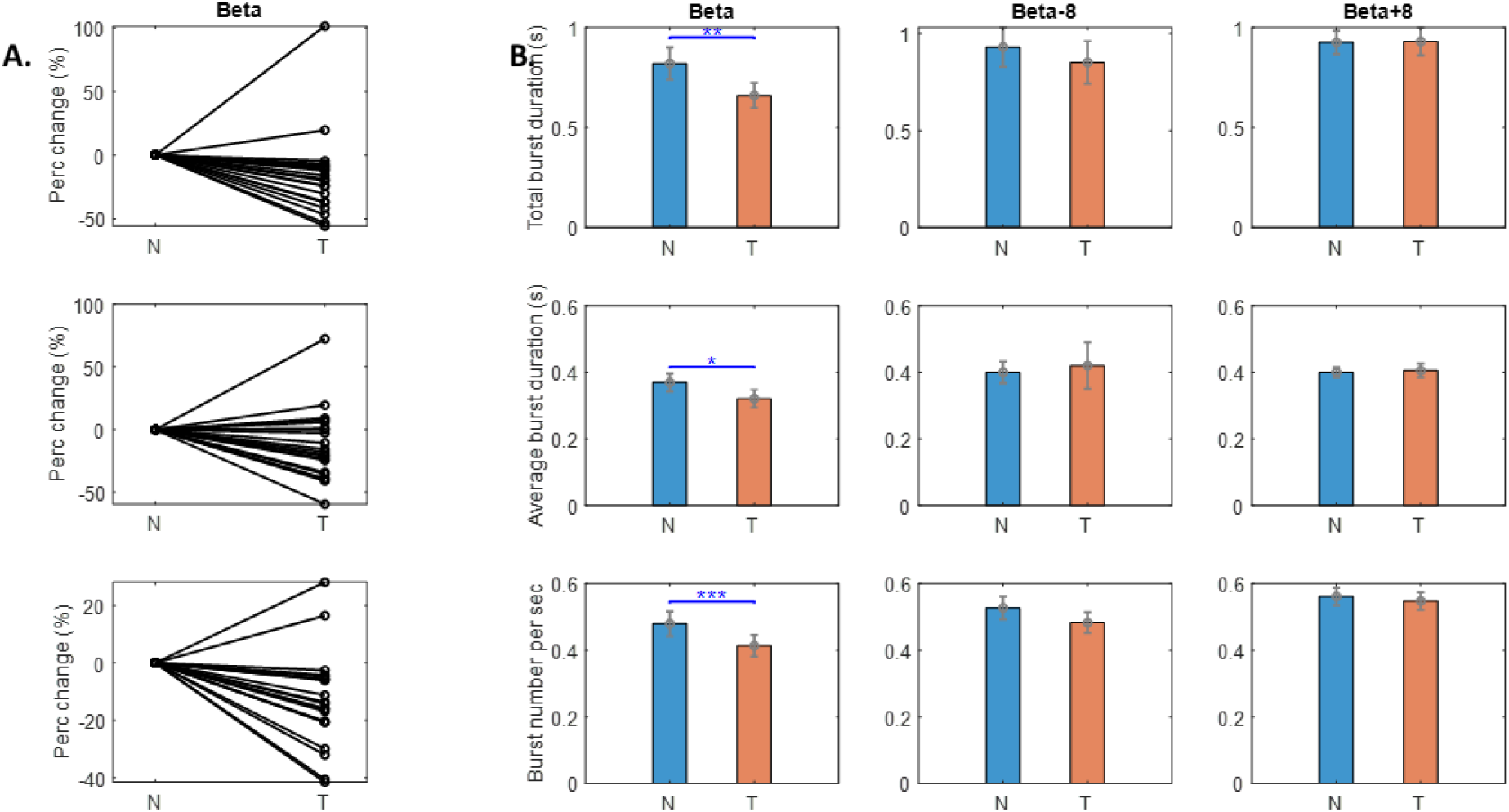
Burst characteristics in LFP changed with neurofeedback training. (A) The percentage change in the total (upper), average (middle), and number of beta bursts for each individual hemisphere. (B) Group average burst characteristics in beta (left), beta-8 (middle), and beta+8 (right) frequency bands in training (T) and no-training (N) conditions. Values are quantified based on the first 4 s data when the ball was shown and presented as mean ± SEM; *p<0.05, **p<0.01, ***p<0.001; Beta indicates person specific beta band.

In order to evaluate whether there was any sustained effect of neurofeedback training, the average beta power was also quantified during the short time window (2 s) when the visual feedback was no longer available, and a black screen was presented before the Go cue. The average beta power during this short time window was also reduced in the training condition compared to the no training condition (k = −0.287 ± 0.103, p = 0.005), and was correlated with the average beta power during the time when the neurofeedback was available (k = 0.23 ± 0.027, p < 0.0001).

### Neurofeedback training reduced beta band synchrony between the conditioned STN and ipsilateral motor cortex

The reduction in beta activity associated with neurofeedback training was not limited to the STN LFP. Beta band activity over the ipsilateral motor cortex in terms of averaged EEG beta power was also reduced during the neurofeedback phase when comparing the training vs no-training condition (F_1,20_ = 9.01, p = 0.0071), even though beta burst characteristics in the EEG were not consistently modulated by the STN LFP based neurofeedback training (F_1,20_ = 1.93, p = 0.1803 for the total duration when beta bursts were detected and F_1,20_ = 1.87, p = 0.1862 for the average duration of each beta burst). In addition, the STN LFP beta targeted neurofeedback training was associated with reduced synchrony between the conditioned STN and the ipsilateral motor cortex in the targeted beta frequency band, as quantified by the phase synchrony index (*PIS*) and spectral coherence (shown in Fig. 6). On average, the *PSI* (F_1,20_ = 14.69, p = 0.001) and spectral coherence (F_1,20_ = 16.81, p = 0.0006) were significantly reduced in the training condition compared with the no-training condition, and this change did not happen in the frequency bands “Beta-8” and “Beta+8” (Fig. 6B).

**Figure 6.**
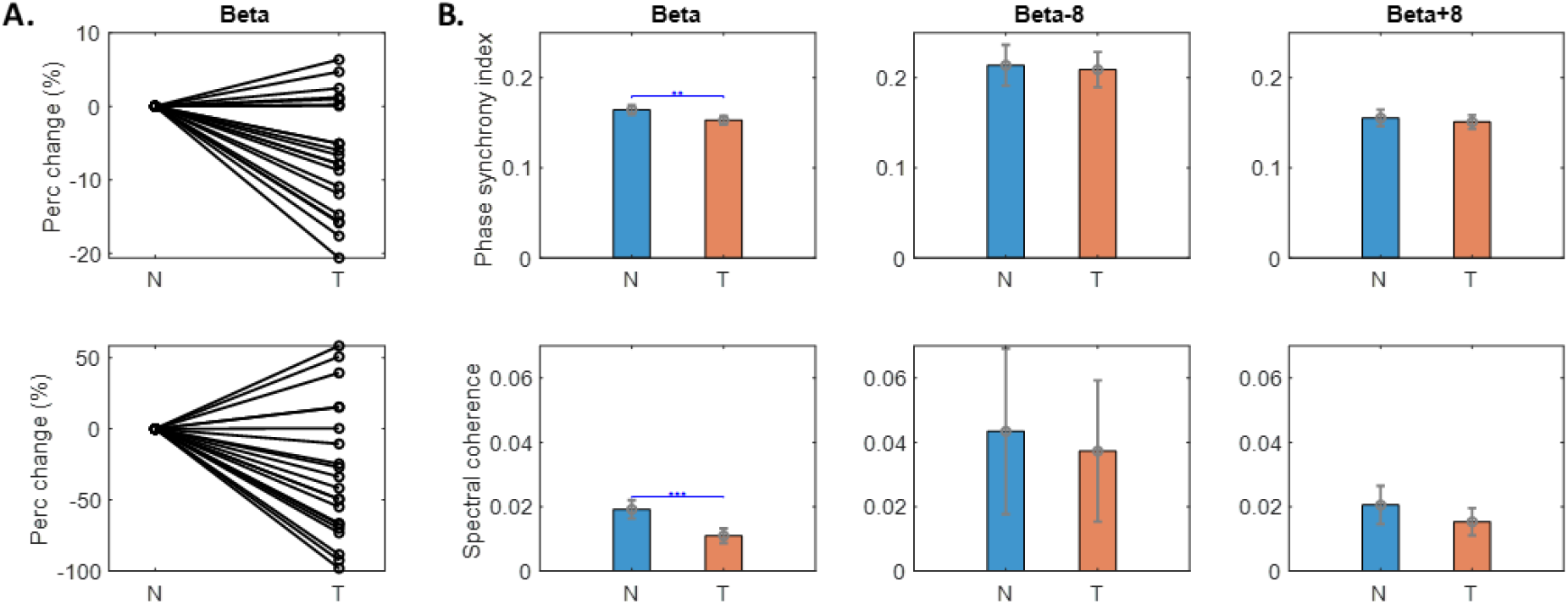
Phase synchrony index and spectral coherence between targeted LFP and ipsilateral EEG was reduced in the beta frequency band with neurofeedback training. (A) The percentage change in the phase synchrony index (upper) and spectral coherence (lower) for each individual hemisphere. (B) Group average phase synchrony index and spectral coherence in beta (left), beta-8 (middle), and beta+8 (right) frequency bands in training (T) and no-training (N) conditions. Values are quantified based on the first 4 s data when the ball was shown and presented as mean ± SEM; **p<0.01; Beta indicates person specific beta band.

### Neurofeedback training speeds up the reaction time in subsequently cued movements

The reaction time in response to the Go cue was significantly reduced in the neurofeedback training condition compared with the no-training condition (390 ± 22 ms compared to 435 ± 28 ms, shown in Fig. 7A), as confirmed by a significant main effect of experimental condition in the two-way ANOVA (F_1,20_ = 22.882, p < 0.001). Since previous analysis of the basketball movements indicated different neurofeedback control performance in different sessions, we examined whether the change in reaction time with neurofeedback training was also session specific. Paired t-tests showed that the reduction of reaction time in the neurofeedback condition was only significant in Session 3 after correcting for multiple comparisons (t_20_ = - 3.289, p = 0.004). This was consistent with the previous results showing best neurofeedback control performance in Session 3. The reaction time increased in Session 4 (443 ± 36 ms) compared to Session 3 (368 ± 28 ms, t_20_ = 3.626, p = 0.002) in the neurofeedback condition; but the reaction time in the no training condition remained constant across sessions (443 ± 30 ms in Session 4 compared to 446 ± 38 ms in Session 3, t_20_ = 0.113, p = 0.911). These results suggest that performance in the participants might have fallen off in the neurofeedback training in Session 4 but remained good for the motor task. There was no consistent improvement in the peak force rate in the pinch movement with the neurofeedback training.

**Figure 7.**
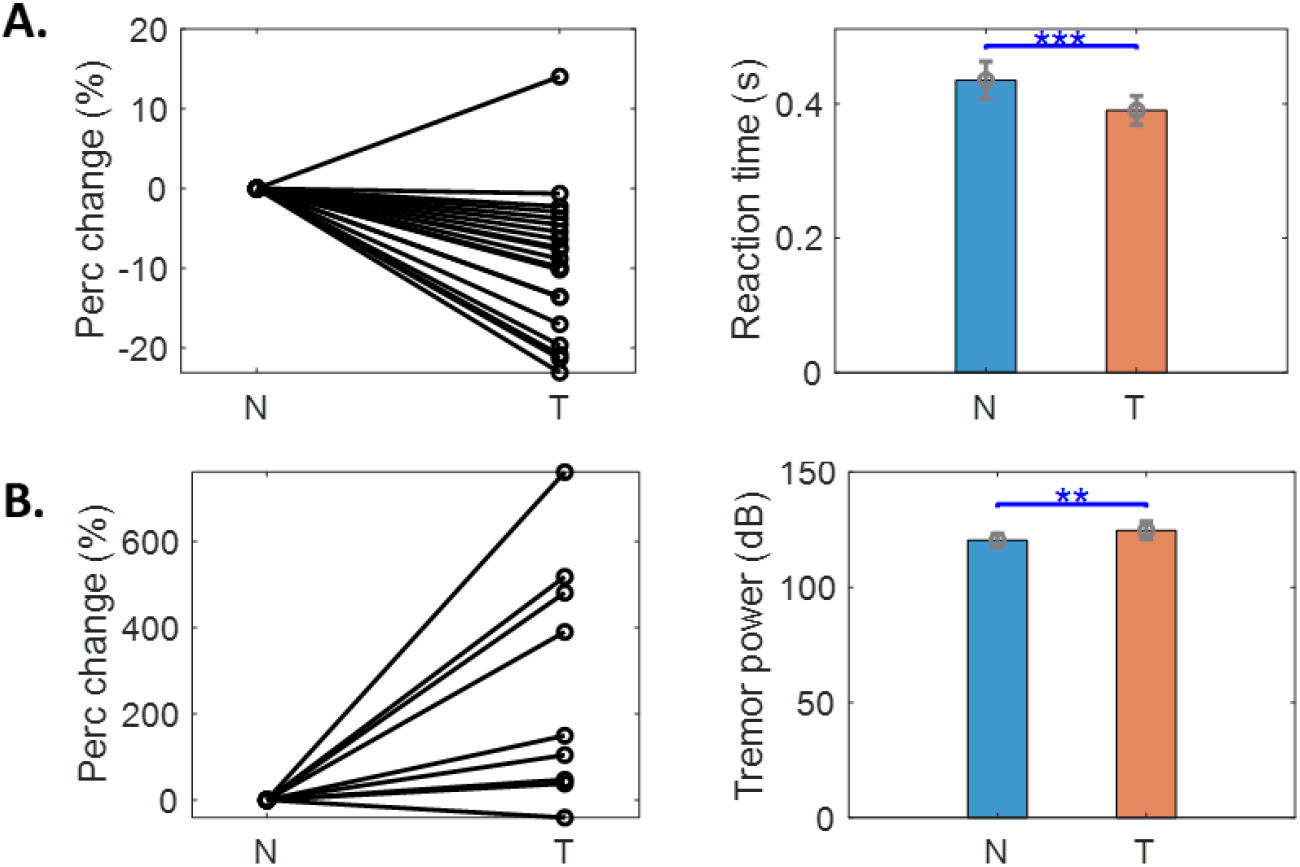
Behavioural changes (reaction time and tremor) associated with neurofeedback training. (A) The percentage change in the reaction time (left) and the group averaged reaction time (right) in training and no-training conditions. Each black line indicates one tested hemisphere on the left. (B) The percentage change in the upper limb tremor tremor power (left) and the group averaged tremor power (right). The tremor power was quantified in 3-7 Hz using the data recorded from tri-axial accelerometers in four patients (8 hemispheres) with tremor during the experiment. Values are presented as mean ± SEM; **p<0.01, ***p<0.001.

Linear mixed effects modelling was used to investigate the relationship between the reaction time and STN LFP activities in the beta (*β*) frequency bands and the experimental condition. Since there is a trend of co-variation in the alpha (α) frequency band, which might also have had an impact on reaction time, we also included the power in the alpha frequency (8-12 Hz) band in the model: *RT* ∼ *k*_1_ ∗ *Training* + *k*_2_ ∗ *β* + *k*_3_ ∗ *α* + 1|*SubID*. This model confirmed the significant effect of beta-targeted neurofeedback training in reducing reaction time (*k*_1_= −0.036 ± 0. 009, p < 0.0001), and of a significant effect of the beta band power 500 ms around the Go cue (*k*_2_= 0.0063 ± 0.0016, p = 0.0001). The positive sign of *k*_2_ indicates reduced reaction time predicted by reduced STN beta band power 500 ms around the Go cue. In comparison, there was no significant effect of the alpha band activity on the RT (*k*_3_= 0.0018 ± 0.0013, p = 0.1697).

### Neurofeedback training targeting beta activity increased tremor

Five out of the twelve participants (one patient had unilateral tremor so rendering 9 STN hemispheres) in the study suffered from tremor and displayed tremor during the recording, which enabled us to investigate how volitional suppression of STN beta oscillations affects tremor in PD. We quantified the severity of the tremor based on the measurements from tri-axial accelerometers attached to the hand. The power of the tremor frequency band (3-7 Hz) in the acceleration in both training and no-training conditions for these 9 hemispheres is shown in Fig. 7B. We found in 5 out of 9 cases that tremor power in the contralateral hand significantly increased during training compared to the no-training condition. Linear mixed effects modelling was used to investigate the relationship between tremor present during the neurofeedback phase and the experimental condition, and STN LFP activities in the beta (β) and tremor (θ) frequency bands (*Tremor* ∼ *k*_1_ ∗ *Training* + *k*_2_ ∗ *β* + *k*_3_ ∗ *θ* + 1|*SubID*). This model confirmed the significant effect of beta-targeted neurofeedback training (*k*_1_= 5.135 ± 0.477, p < 0.0001) on increasing tremor, and also indicated that increased tremor band activity in the STN LFP (*k*_3_= 0.863 ± 0.043, p < 0.0001), together with reduced beta band activity in the STN LFP (*k*_2_= −0.326 ± 0.119, p = 0.006) were associated with tremor.

### Cross day learning effect of the neurofeedback training

In most EEG based neurofeedback studies training sessions have been conducted over several separate days (Engelbregt et al. 2016, Schabus et al. 2017). In this study, 4 participants (8 hemispheres) repeated the task on two separate, consecutive days. On Day 2, all the 8 tested hemispheres showed significant difference in the ball’s final positions between the training and no-training conditions (considering all individual trials using non-paired t-test for different hemispheres separately). Comparing against Day 1, 6 out of the 8 tested hemispheres showed increased neurofeedback control (difference in the training and no-training condition) on Day 2 (shown in Fig. 8A). The other 2 tested hemispheres which had already achieved good neurofeedback control on Day 1 did not further improve on Day 2 (H7 and H8 in Fig. 8A).

**Figure 8.**
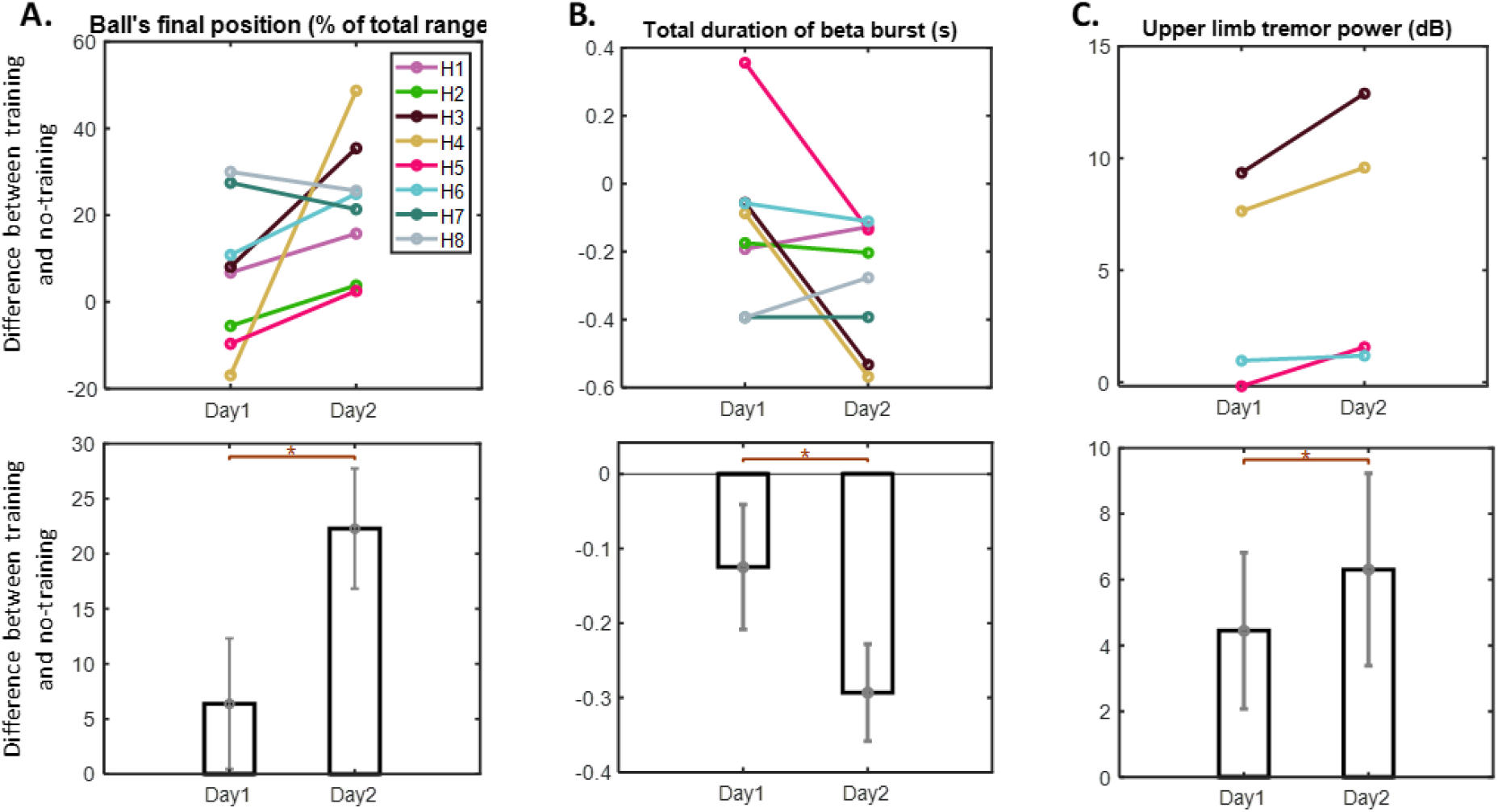
Comparison between two training days. (A) The difference in the ball’s final position between training and no-training conditions on Day1 and Day2. (B) The difference in the total burst duration between training and no-training conditions on Day1 and Day2. (C) The difference in physical tremor power between training and no-training conditions on Day1 and Day2. Individual hemispheres and group average data are shown in the upper and lower panels, respectively. Values are presented as mean ± SEM; *p<0.05 (Wilcoxon signed rank test).

LME modelling confirmed significant interaction between experimental condition and recording days on the total duration of beta bursts (k = −163.94 ± 65.579, p = 0.013). As shown in Fig. 8B, compared to Day 1, the neurofeedback training on Day 2 was associated with more reduction in the total beta burst duration compared to the no-training condition. These results suggested that a better training effect was achieved on Day 2 compared to Day 1. LME modelling also showed that the reaction time was consistently reduced during Day 2 compared to Day 1 in the training condition only (k = −0.026 ± 0.011, p = 0.014); whereas cross-day difference in the no-training condition was not significant (k = −0.024 ± 0.016, p = 0.139). For all the individual trials across the two recording days, the strongest predictor for the reaction time to the cued movements was still the beta power in the STN LFP 500 ms around the Go cue (k = 0.0057 ± 0.0010, p< 0.001).

For the 2 patients (4 hemispheres) who had tremor and repeated the task over two consecutive days, tremor during the training condition was increased more on Day 2 than Day 1 in all four hemispheres (Fig. 8C). Considering all the individual trials across the two recording days, the strongest predictor for the tremor was the tremor band power (3-7 Hz) in the STN LFP (k = 0.694 ± 0.050, p< 0.001).

## Discussion

In this study, we tested a sequential neurofeedback-behaviour task paradigm in which patients with Parkinson’s disease learned to self-suppress pathological beta bursts in the STN before a cued finger movement. The online visual feedback provided during the neurofeedback phase indexed whether STN beta power over the preceding 500ms exceeded the 75^th^ percentile, and therefore was also related to the incidence and duration of beta bursts during this time window. This experimental design afforded a relatively smooth visual feedback (with vertical position fixed between updates at 250ms intervals) and allowed patients to gain a sense of agency when controlling the visual feedback within 30 mins. Training with this paradigm led to reduced incidence of beta bursts and reduced average burst duration in the STN LFP, and reduced coherence between the STN and ipsilateral motor cortex in the targeted beta frequency band. Importantly, this is the first study to show that volitional suppression of beta bursts in the STN LFP facilitated by neurofeedback training is able to speed up movement initialisation in a cued movement in PD patients. This is consistent with previous studies which demonstrated a positive correlation between volitionally induced beta-power and reaction time, but not between beta power and movement velocity-parameters (Khanna and Carmena, 2017; Peles et al., 2019). However, a lack of effect on the rate of force development in this study may also relate to the experimental design. In this study, the patients were required to pinch as fast as they could to reach a predefined comfortable force level. The target force level was only 50 % of their maximum force. In addition, the feedback provided to the participants about the pinch movement after each movement was determined by the reaction time only, therefore only performance of this measure was reinforced during the paradigm.

### Neurofeedback training for PD

Neurofeedback, proposed as a voluntary operant conditional training for self-regulation of brain activities, has been investigated in the treatment of epilepsy, anxiety, depression, and attention deficit/hyperactivity disorder (ADHD). Despite questions over its scientific rigour and clinical relevance (Schabus et al. 2017, Thibault et al. 2017, Thibault and Raza, 2016), there is increasing interest in exploring the use of neurofeedback as a therapeutic technique for Parkinson’s disease (Esmail and Linden 2014, Carney 2019). For example, functional MRI neurofeedback guided motor imagery has been used to upregulate activity in the supplementary motor area (SMA) in patients with PD, and this was concomitant with improvement in the motor scale of the Unified PD Rating Scale (Subramanian et al. 2011). However, a later study showed that motor improvement was not significantly different from a control group who experienced similar motor training without veritable neurofeedback (Subramanian et al. 2016).

Cortical sensorimotor rhythms measured using EEG have also been used as the target for neurofeedback training in Parkinson’s disease. However, effects on motor behaviour have been contradictory. Philippens et al. (2017) reported that increasing the sensorimotor rhythm between 12–17 Hz in the sensorimotor cortex through neurofeedback training in parkinsonian nonhuman primates reduced 1-methyl-4-phenyl-1,2,3,6-tetrahydropyridine (MPTP) - induced parkinsonian signs such as immobility, muscle rigidity, rest tremor, apathy and inadequate grooming. On the other hand, Khanna and Carmena (2017) showed that reaching movements preceded by a reduction in beta power achieved through neurofeedback training exhibited significantly faster movement onset times than those preceded by an increase in beta power in healthy macaque monkeys. Similarly, both healthy human participants and patients with Parkinson’s disease can learn to voluntarily increase or decrease oscillations in the beta frequency band measured over sensorimotor cortex after neurofeedback training. Boulay et al. (2011) reported that increasing sensorimotor beta rhythms in healthy human participants was associated with longer reaction times than decreasing sensorimotor beta rhythms. In particular, pre-movement sensorimotor beta amplitude was correlated with motor performance in subjects with initial poor performance: lower amplitude was associated with faster and more accurate movement (McFarland et al., 2015). However, training PD patients to up-regulate oscillatory activity in the 8–15 Hz range (which falls within the low beta range) and down-regulate activity in the 23–34 Hz (high beta) failed to induce any significant clinical effect (Erickson-Davis et al. 2012).

In the current study, the online feedback was provided based on the beta band activity in the STN LFP signals recorded from the electrode implanted for deep brain stimulation. This activity has been previously shown to be related to motor impairments in PD (Jenkinson and Brown, 2011). We selected a patient-specific beta frequency band which was modulated by voluntary movements and was also enhanced relative to other frequency bands during rest (displayed as a peak in the power spectra). We showed that volitional suppression of bursts of this activity was accompanied by reduced reaction time in cued movements.

Another important observation in this study is that neurofeedback training targeting beta oscillations may increase tremor, as well as tremor band activities in the STN LFPs in tremulous patients. Our results show that increased tremor band activity and reduced beta band activity in the STN LFP predict more severe tremor during the neurofeedback phase in these patients. This is consistent with previous studies showing that in the presence of tremor, neuronal oscillations at tremor frequency (3–7 Hz) tend to increase in the cortical-basal ganglia-thalamic circuit (Hirschmann et al., 2013); whereas beta power (13–30 Hz) and beta band coupling in the motor network are reduced (Qasim et al., 2016). It is possible that disengaging neuronal activity from the beta band may allow synchronisation in the tremor frequency band, and lead to increased tremor. Therefore, neurofeedback training targeting reduction in beta activity might not help patients with tremor. Such patients might be better served by neurofeedback training focussing on tremor related oscillations.

### fMRI vs EEG/LFP signals for neurofeedback training

Real-time functional MRI (rtfMRI) and EEG are the most often used imaging modalities for neurofeedback training. It has been noted that EEG neurofeedback requires many sessions of training for a participant to alter electrical activity, compared with rtfMRI in which participants can selectively modify the fMRI blood-oxygen-level dependent (BOLD) signal within 30 min of training (Thibault et al 2016). This is despite the much longer time delay between a change in neural activity and its reflection in the BOLD signal (and thus on the feedback display). For example, in the study of Subramanian et al. (2011, 2016), patients were informed that the delay was approximately 5 s, and yet patients were able to upregulate the BOLD activity of sensorimotor cortex within 2 sessions. In contrast, it takes 6-14 sessions (Boulay et al, 2011; McFarland 2015) or even more (24 training sessions in Erickson-Davis et al. (2012)) for participants to be able to modulate oscillatory activities in the beta frequency band. This may be due to the lower signal to noise ratio in the EEG/LFP measurements and the fast time dynamics in the signal of interest. Synchrony in the beta frequency band in the motor cortex or in the basal ganglia takes the form of bursts of different durations and amplitudes (2015; Tinkhauser et al., 2017a, b). Directly translating continuously updated beta-band power into visual feedback can lead to a flickering, dynamic feedback which can be confusing for participants. Limiting the update rate, averaging data over a time window of seconds in duration, and converting continuous measurements of power into a binary visual display are often used strategies to smooth the feedback signal in EEG neurofeedback. In our paradigm, we calculated the beta band power within a 500 ms window from STN LFPs and updated the visual feedback every 250 ms. This was based on previous studies showing that long beta bursts (over 500 ms) are more closely related to the symptoms of bradykinesia and rigidity in Parkinson’s disease. By taking into account the temporal dynamics of the signal of interest, and reducing the variance and noise in the visual feedback that are less related to the underlying pathology, our paradigm allowed patients to learn to suppress beta power within 30 min of training.

Our results also showed that there was concurrent reduction in beta band activities over the motor cortex measured using EEG, and reduced coherence in the STN-M1 motor network in the beta frequency band. This raises the question would it be equally effective to use EEG based neurofeedback targeting the beta bursts over sensorimotor cortex measured using EEG? This was not directly tested here, but the temporal dynamics of the beta oscillation in the STN that were used to provide feedback, and which were modulated by feedback, failed to be modulated in the EEG. Moreover, in those patients undergoing surgery for deep brain stimulation, neurofeedback training based on LFP signals recorded from the stimulation electrodes could be easily implemented, given the recent technical advance that allows chronic sensing of LFPs and wireless transmission of data to a hand held device (Stanslaski et al., 2012). In theory, such patients could use neurofeedback training, in addition to the effects of stimulation, to acquire mental strategies that improve motor performance.

## Limitations

There are some limitations in the current study. First, the study was conducted between surgery for the implantation and externalization of DBS electrodes and surgery for connecting the electrodes to a subcutaneous pulse generator. Thus the time window available to conduct the experiment was very limited in each participant. Our results showed that the performance of the patients in suppressing beta bursts with neurofeedback training improved from Session 1 to Session 3 within Day 1, but the performance deteriorated in Session 4, possibly due to fatigue. The patients who performed the task over two days showed significant improvement in the performance of controlling the visual feedback on Day 2 compared to Day 1. With chronic sensing it will be possible to access the STN LFP as required (Khanna et al., 2017; Herron et al., 2017; Haddock et al., 2018; Houston et al., 2018). It would therefore be interesting to test the effect of neurofeedback training over longer periods when bidirectional devices become more widely available. In our study, the ‘test’ motor task used to evaluate the carry over effects of neurofeedback training immediately followed neurofeedback. We did not test whether there was any long-term carry over effect of the neurofeedback training on motor symptoms associated with Parkinson’s disease. Finally, we did not use ‘sham-feedback’ as a control condition. However, we suspected that intermixing ‘sham-feedback’ and ‘veritable feedback’ might have had negative impact on motivation and might have interfered with learning. But without a sham-feedback condition we cannot exclude the possibility that the mental strategy adopted was alone sufficient to suppress beta and change performance without feedback. The effect of attention alone was controlled for in our no-training condition.

In summary, neurofeedback based on STN beta power led to volitional suppression of beta activity, and particularly of beta bursts in the STN, and a reduction in the coupling of STN beta activity with EEG over the motor cortex. These changes were accompanied by a reduction in cued reaction time. The results strengthen the link between subthalamic beta oscillations, and beta bursts in particular, and motor impairment. However, although the effects of neurofeedback on motor initialisation are encouraging it remains to be seen if there is more prolonged carry-over of voluntary control, and whether this translates into clinically meaningful symptom amelioration.

## ACKNOWLEDGEMENTS

This work was supported by the MRC (MR/P012272/1 and MC_UU_12024/1) and the Rosetrees Trust.

## References

Azarpaikan A, Torbati HT, Sohrabi M (2014). Neurofeedback and physical balance in Parkinson’s patients. Gait & posture, 40(1): 177–81.

Boulay CB, Sarnacki WA, Wolpaw JR, McFarland DJ (2011). Trained modulation of sensorimotor rhythms can affect reaction time. Clinical Neurophysiology, 122(9): 1820–6.

Carney RS (2019). Neurofeedback Training Enables Voluntary Alteration of β-Band Power in the Subthalamic Nucleus of Individuals with Parkinson’s Disease. eNeuro, 6(2).

de Hemptinne C, Ryapolova-Webb ES, Air EL, Garcia PA, Miller KJ, Ojemann JG, Ostrem JL, Galifianakis NB, Starr PA (2013). Exaggerated phase–amplitude coupling in the primary motor cortex in Parkinson disease. Proceedings of the National Academy of Sciences of the United States of America, 110(12): 4780–5.

de Hemptinne C, Swann NC, Ostrem JL, Ryapolova-Webb ES, Luciano MS, Galifianakis NB, Starr PA (2015). Therapeutic deep brain stimulation reduces cortical phase-amplitude coupling in Parkinson’s disease. Nature Neuroscience, 18: 779–86.

Engelbregt HJ, Keeser D, Van Eijk L, Suiker EM, Eichhorn D, Karch S, Deijen JB, Pogarell O (2016). Short and long-term effects of sham-controlled prefrontal EEG-neurofeedback training in healthy subjects. Clinical Neurophysiology, 127(4): 1931–7.

Erickson-Davis CR, Anderson JS, Wielinski CL, Richter SA, Parashos SA (2012). Evaluation of neurofeedback training in the treatment of Parkinson’s disease: A Pilot Study. Journal of Neurotherapy, 16(1): 4–11.

Esmail, S and Linden, DE (2014). Neural networks and neurofeedback in Parkinson’s disease. Neuroregulation, 1(3-4): 240.

Fischer P, Pogosyan A, Cheeran B, Green AL, Aziz TZ, Hyam J, Little S, Foltynie T, Limousin P, Zrinzo L, Hariz M, Samuel M, Ashkan K, Brown P, Tan H (2017). Subthalamic nucleus beta and gamma activity is modulated depending on the level of imagined grip force. Experimental Neurology, 293: 53–61.

Fukuma R, Yanagisawa T, Tanaka M, Yoshida F, Hosomi K, Oshino S, Tani N, Kishima H (2018). Real-time neurofeedback to modulate β-band power in the subthalamic nucleus in Parkinson’s disease patients. eNeuro, 5(6).

George JS, Strunk J, Mak-McCully R, Houser M, Poizner H, Aron AR (2013). Dopaminergic therapy in Parkinson’s disease decreases cortical beta band coherence in the resting state and increases cortical beta band power during executive control. NeuroImage: Clinical, 3: 261–70.

Haddock A, Mitchell KT, Miller A, Ostrem JL, Chizeck HJ, Miocinovic S (2018). Automated deep brain stimulation programming for tremor. IEEE Transactions on Neural Systems and Rehabilitation Engineering, 26(8): 1618–25.

Herron JA, Thompson MC, Brown T, Chizeck HJ, Ojemann JG, Ko AL (2017). Chronic electrocorticography for sensing movement intention and closed-loop deep brain stimulation with wearable sensors in an essential tremor patient. Journal of neurosurgery, 127(3): 580–7.

Hirschmann J, Hartmann CJ, Butz M, Hoogenboom N, Özkurt TE, Elben S, Vesper J, Wojtecki L, Schnitzler A (2013). A direct relationship between oscillatory subthalamic nucleus–cortex coupling and rest tremor in Parkinson’s disease. Brain, 136(12): 3659–70.

Houston B, Thompson M, Ko A, Chizeck H (2018). A machine-learning approach to volitional control of a closed-loop deep brain stimulation system. Journal of neural engineering, 16(1): 016004.

Jenkinson N and Brown P (2011). New insights into the relationship between dopamine, beta oscillations and motor function. Trends in Neurosciences, 34: 611–8.

Kato K., Yokochi F., Taniguchi M., Okiyama R., Kawasaki T., Kimura K., Ushiba Junichi (2015). Bilateral coherence between motor cortices and subthalamic nuclei in patients with Parkinson’s disease. Clinical Neurophysiology, 126: 1941–50

Khanna P, Swann NC, de Hemptinne C, Miocinovic S, Miller A, Starr PA, Carmena JM (2017). Neurofeedback control in parkinsonian patients using electrocorticography signals accessed wirelessly with a chronic, fully implanted device. IEEE Transactions on Neural Systems and Rehabilitation Engineering, 25(10): 1715–24.

Khanna P and Carmena JM (2017). Beta band oscillations in motor cortex reflect neural population signals that delay movement onset. Elife, 6: e24573.

Kühn AA, Kupsch A, Schneider GH, Brown P (2006). Reduction in subthalamic 8-35 Hz oscillatory activity correlates with clinical improvement in Parkinson’s disease. European Journal of Neuroscience, 23 (7): 1956–60.

Kühn AA, Tsui A, Aziz T, Ray N, Brücke C, Kupsch A, Schneider GH, Brown P (2009). Pathological synchronisation in the subthalamic nucleus of patients with Parkinson’s disease relates to both bradykinesia and rigidity, Experimental neurology, 215(2): 380–7.

Lachaux JP, Rodriguez E, Martinerie J, Varela FJ (1999). Measuring phase synchrony in brain signals. Human brain mapping, 8(4): 194–208.

Lachaux JP, Rodriguez E, Le Van Quyen M, Lutz A, Martinerie J, Varela FJ (2000). Studying single-trials of phase synchronous activity in the brain. International Journal of Bifurcation and Chaos, 10(10): 2429–39.

Little S, Pogosyan A, Kuhn AA, Brown P (2012). Beta band stability over time correlates with Parkinsonian rigidity and bradykinesia, Experimental neurology, 236(2): 383–8.

Little S, Pogosyan A, Neal S, Zavala B, Zrinzo L, Hariz M, Foltynie T, Limousin P, Ashkan K, FitzGerald J, Green AL, Aziz TZ, Brown P (2013). Adaptive deep brain stimulation in advanced Parkinson disease. Annals of neurology, 74 (3): 449–57.

Little S, Beudel M, Zrinzo L, Foltynie T, Limousin P, Hariz M, Neal S, Cheeran B, Cagnan, H, Gratwicke J, Aziz TZ, Pogosyan A, Brown P (2016). Bilateral adaptive deep brain stimulation is effective in Parkinson’s disease. Journal of Neurology, Neurosurgery, and Psychiatry, 87(7): 717–21.

McFarland DJ, Sarnacki WA, Wolpaw JR (2015). Effects of training pre-movement sensorimotor rhythms on behavioral performance. Journal of neural engineering, 12(6): 066021.

Peles O, Werner-Reiss U, Bergman H, Israel Z, Vaadia E (2019). Phase-specific microstimulation in brain-machine interface setting differentially modulates beta oscillations and affects behavior. bioRxiv, 622787.

Philippens IH, Wubben JA, Vanwersch RA, Estevao DL, Tass PA (2017). Sensorimotor rhythm neurofeedback as adjunct therapy for Parkinson’s disease. Annals of clinical and translational neurology, 4(8): 585–90.

Qasim SE, de Hemptinne C, Swann NC, Miocinovic S, Ostrem JL, Starr PA (2016). Electrocorticography reveals beta desynchronization in the basal ganglia-cortical loop during rest tremor in Parkinson’s disease. Neurobiology of disease, 86: 177–86.

Ros T, Baars BJ, Lanius RA, Vuilleumier P (2014). Tuning pathological brain oscillations with neurofeedback: a systems neuroscience framework. Frontiers in Human Neuroscience, 8: 1008.

Rowland NC, de Hemptinne C, Swann NC, Qasim S, Miocinovic S, Ostrem JL, Knight RT, Starr PA (2015). Task-related activity in sensorimotor cortex in Parkinson’s disease and essential tremor: Changes in beta and gamma bands. Frontiers in Human Neuroscience, 9: 512.

Schabus M, Griessenberger H, Gnjezda MT, Heib DP, Wislowska M, Hoedlmoser K (2017). Better than sham? A double-blind placebo-controlled neurofeedback study in primary insomnia. Brain, 140(4): 1041–52.

Schneider SA, Edwards MJ, Mir P, Cordivari C, Hooker J, Dickson J, Quinn N, Bhatia KP (2007). Patients with adult-onset dystonic tremor resembling parkinsonian tremor have scans without evidence of dopaminergic deficit (SWEDDs). Movement disorders: official journal of the Movement Disorder Society, 22(15): 2210–5.

Stanslaski S, Afshar P, Cong P, Giftakis J, Stypulkowski P, Carlson D, Linde D, Ullestad D, Avestruz AT, Denison T (2012). Design and validation of a fully implantable, chronic, closed-loop neuromodulation device with concurrent sensing and stimulation. IEEE Transactions on Neural Systems and Rehabilitation Engineering, 20(4): 410–21.

Subramanian L, Hindle JV, Johnston S, Roberts MV, Husain M, Goebel R, Linden D (2011). Real-time functional magnetic resonance imaging neurofeedback for treatment of Parkinson’s disease. Journal of Neuroscience, 31(45): 16309–17.

Subramanian L, Morris MB, Brosnan M, Turner DL, Morris HR, Linden DE (2016). Functional magnetic resonance imaging neurofeedback-guided motor imagery training and motor training for Parkinson’s disease: randomized trial. Frontiers in behavioral neuroscience, 10: 111.

Tan H, Pogosyan A, Ashkan K, Green AL, Aziz T, Foltynie T, Limousin P, Zrinzo L, Hariz M, Brown P (2016). Decoding gripping force based on local field potentials recorded from subthalamic nucleus in humans. Elife, 5: e19089.

Thibault RT, Lifshitz M, Raz A (2016). The self-regulating brain and neurofeedback: experimental science and clinical promise. Cortex, 74: 247–61.

Thibault RT and Raz A (2016). When can neurofeedback join the clinical armamentarium? Lancet, 3(6).

Thibault RT, Lifshitz M, Raz A (2017). The climate of neurofeedback: scientific rigour and the perils of ideology. Brain, 141(2): e11.

Tinkhauser G, Pogosyan A, Little S, Beudel M, Herz DM, Tan H, Brown P (2017a). The modulatory effect of adaptive deep brain stimulation on beta bursts in Parkinson’s disease. Brain, 140 (4): 1053–67.

Tinkhauser G, Pogosyan A, Tan H, Herz DM, Kühn AA, Brown P (2017b). Beta burst dynamics in Parkinson’s disease OFF and ON dopaminergic medication. Brain, 140 (11): 2968–81.

Tinkhauser G, Torrecillos F, Duclos Y, Tan H, Pogosyan A, Fischer P, Carron R, Welter ML, Karachi C, Vandenberghe W, Nuttin B, Witjas T, Régis J, Azulay JP, Eusebio A, Brown P (2018). Beta burst coupling across the motor circuit in Parkinson’s disease. Neurobiology of Disease, 117: 217–25.

